# Site Frequency Spectrum of the Bolthausen-Sznitman Coalescent

**DOI:** 10.1101/799627

**Authors:** Götz Kersting, Arno Siri-Jégousse, Alejandro H. Wences

**Affiliations:** Goethe Universität, Institut für Mathematik, Frankfurt am Main, Germany; UNAM, IIMAS, Departamento de Probabilidad y Estadística, Mexico

**Keywords:** Bolthausen-Sznitman coalescent, Site frequency spectrum, Random recursive tree, Branch lengths

## Abstract

We derive explicit formulas for the two first moments of he site frequency spectrum (*SFS*_*n,b*_)_1≤*b*≤*n*−1_ of the Bolthausen-Sznitman coalescent along with some precise and efficient approximations, even for small sample sizes *n*. These results provide new *L*_2_-asymptotics for some values of *b* = *o*(*n*). We also study the length of internal branches carrying *b* > *n/*2 individuals. In this case we obtain the distribution function and a convergence in law. Our results rely on the random recursive tree construction of the Bolthausen-Sznitman coalescent.

## 1 Introduction

The Bolthausen-Sznitman coalescent is an exchangeable coalescent with multiple collisions that has recently gained attention in the theoretical population genetics literature. It has been described as the limit process of the genealogies of different population evolution models, including models that contemplate the effect of natural selection [15, 16]. It has also been proposed as a new null model for the genealogies of rapidly adapting populations, such as pathogen microbial populations, and other populations that show departures from King-man’s null model [1, 13].

A measure of the genetic diversity in a present day sample of a population is often used in population genetics in order to infer its evolutionary past and the forces at play in its dynamics. The *Site Frequency Spectrum* (SFS) is a well known theoretical model of the genetic diversity present in a population, it assumes that neutral mutations arrive to the population as a Poisson Process and that each arriving mutation falls in a different site of the genome (infinite sites model), in contrast to the *Allele Frequency Spectrum* in which mutations are assumed to fall on the same site but create a new allele every time (infinite alleles model). Given the close relation between the Site Frequency Spectrum and the whole structure of the underlying genealogical tree, it can be used as a model selection tool for the evolutionary dynamics of a population [3, 10, 4].

In this work we give explicit expressions of the first and second moments for the whole Site Frequency Spectrum (*SFS*_*n,b*_)_1≤*b*<*n*_ of the Bolthausen-Sznitman coalescent, which to our knowledge were only known for Kingman’s coalescent until now [5]. Here *SFS*_*n,b*_ denotes the number of mutations shared by *b* individuals in the sample of size *n*. For the expectation we obtain the formula

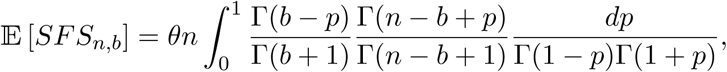

where *θ* denotes the mutation rate. For larger values of *n* there might occur problems in the calculation of this integral due to the exorbitant growth of the Gamma function. Also this formula allows no insight into the shape of the expected site frequency spectrum. For this purpose approximations are helpful. A first approximation, resting on Stirling’s formula, reads for 2 ≤ *b* ≤ *n* – 1

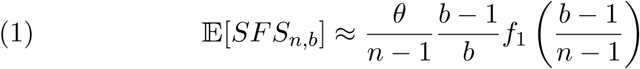

where *f*_1_ is a convex, non-monotone function on (0, 1) defined by

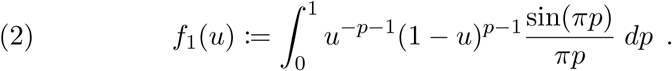

We remark that this integral may be reduced to the (complex) exponential integral *Ei*(·). These formulas show that the shape of the Site Frequency Spectrum, restricted to the range 2 ≤ *b* < *n*, is explained essentially by one function not depending on the population size *n*. Also our approximations update those given in [13] for the case of families with frequencies close to 0 and 1, since we have *f*_1_(*u*)∼(*u* log *u*)^−2^ close to 0 and *f*_1_(*u*)∼((*u* − 1) log(1 − *u*))^−1^ close to 1, see equations (30) and (31) below. The case *b* = 1 is not covered by (1), it has to be treated separately, which reflects the dominance of external branches in the Bolthausen-Sznitman coalescent. See Theorem 3.4 for a complete summary.

The above approximation is accurate also from a numerical point of view. Only for *b* = 2 we encounter an enlarged relative error which anyhow remains less than 10 percent for *n* ≥ 8. If a more precise result is desired then the following refined approximation may be applied for 2 ≤ *b* ≤ *n*:

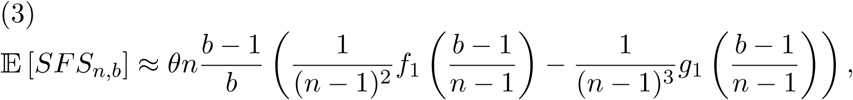

with a positive function *g*_1_ on (0, 1) given by

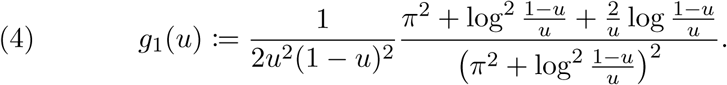

With this formula we have a relative error remaining below 1 percent for *b* = 2 and *n* ≥ 10, below 0.5 percent for *b* = 2 and *n* ≥ 150, and below 0.3 percent for *b* ≥ 3 and *n* ≥ 10. Thus this approximation appears well-suited for practical purpose. Figure 1 illustrates its precision in the cases *n* = 5, 20, 35 and *θ* = 1.

**Figure 1:**
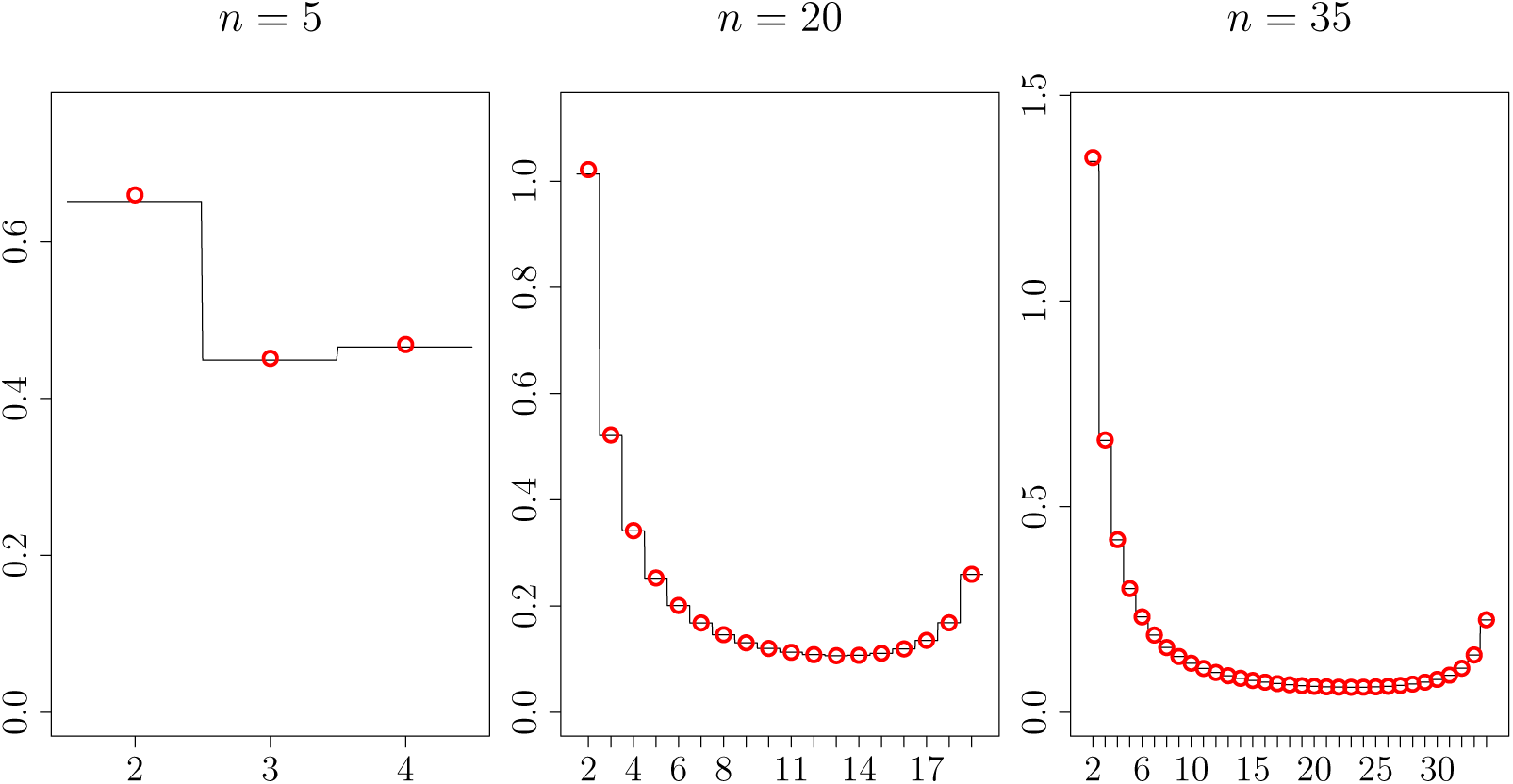
Comparison of exact and approximated values of 𝔼 [*SFS*_*n,b*_], red circles present the exact values for *b* = 2 to *n* − 1, and the black lines their refined approximations (3).

For *b* = 1 the approximation formula corresponding to (1) reads

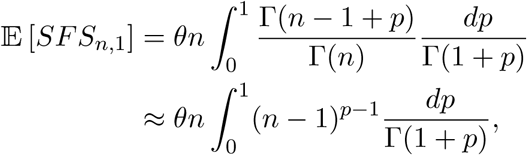

which is an immediate consequence of Stirling’s approximation. It is precise for small *n* and requires no further correction as in the case *b* ≥ 2.

We also study the asymptotic behavior of the second moments which, together with the above asymptotics for the first moment, leads to the following *L*^2^ convergences:

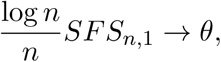

and, whenever *b* ≥ 2 and 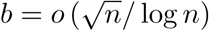,

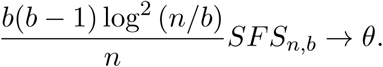

These generalize and strengthen the results in [2] for the Bolthausen-Sznitman coalescent.

We also provide the joint distribution function of the branch lengths of large families, i.e families of size at least half the total population size, and their marginal distribution function. These results are useful to obtain the marginal distribution function of the Site Frequency Spectrum and a sampling formula for the half of the vector corresponding to large family sizes, although we do not present such tedious computations here.

Asymptotic results for related functionals on the Bolthausen-Sznitman coalescent have been derived by studying the block count chain of the coalescent through a coupling with a random walk as in [8] and [9], where asymptotics for the total number of jumps, and the total, internal, and external branch lengths of the Bolthausen-Sznitman coalescent are described; these results give the asymptotic behaviour of the total number of mutations present in the population, the number of mutations present in a single individual, and the number of mutations present in at least 2 individuals. Also, a Markov chain approximation of the initial steps of the process was developed in [2] where asymptotics for the total tree length and the Site Frequency Spectrum of small families were derived for a class of Λ-coalescents containing the Bolthausen-Sznitman coalescnet.

Progress has also been made for the finite coalescent even for the general coalescent process. The finite Bolthausen-Sznitman coalescent has been studied through the spectral decomposition of its jump rate matrix described in [11] where the authors used it to derive explicit expressions for the transition probabilities and the Green’s matrix of this coalescent, and also the Kingman coalescent. The spectral decomposition of the jump rate matrix of a general coalescent, including coalescents with multiple mergers, is also used in [17] where an expression for the expected Site Frequency Spectrum is given in terms of matrix operations which in the case of the Bolthausen-Sznitman coalescent result in an algorithm requiring on the order of *n*^2^ computations. In [7] another expression in terms of matrix operations is given for this and other functionals on general coalescent processes, both in expected value (and higher moments) and in distribution; these expressions however are deduced from the theory of phase-type distributions, in particular distributions of rewards constructed on top of coalescent processes, and also require vast computations for large population sizes.

Our method, mainly based on the Random Recursive Tree construction of the Bolthausen-Sznitman coalescent given in [6], gives easy-to-compute expressions for the first and second moments of the Site Frequency Spectrum of this particular coalescent. This combinatorial construction not only allows us to study the bottom but also the top of the tree thus providing an additional insight into the past of the population and large families, both asymptotically and for any fixed population size.

In Section 2 we layout the basic intuitions that compose the bulk of our method, including the Random Recursive Tree construction of the Bolthausen-Sznitman coalescent and the derivation of the first moment of the Site Frequency Spectrum for the infinite coalescent as a first application (Corollary 2.2). In Section 3 we present our results on the first and second moments of the branch lengths (Theorem 3.1) and of the Site Frequency Spectrum (Corollary 3.3) for any fixed family size and initial population. We then use these expressions to obtain asymptotic approximations of these moments as the initial population goes to infinity (Theorems 3.4 and 3.5) which lead to *L*^2^ convergence results on the SFS (Corollary 3.3). In Section 4 we restrict ourselves to the case of large family sizes and present the joint and marginal distribution functions of their branch lengths (Theorems 4.1 and 4.3), along with a limit in law result (Corollary 4.4). Section 5 provides explanations for approximations (1) and (3). Finally, in Sections 6 and 7 we provide detailed proofs of our results.

## 2 Preliminaries

Consider the Bolthausen-Sznitman coalescent (Π^∞^(*t*))_*t*≥0_ with values in *𝒫*_∞_, the space of partitions of ℕ, and the ranked coalescent (| П^∞^ (*t*)| ^↓^)_*t* ≥0_ with values in the space of mass partitions *𝒫* _[0,1]_, made of the asymptotic frequencies of Π^∞^(*t*) reordered in a non-increasing way. In what follows we present the Random Recursive Tree (RRT) construction of the Bolthausen-Sznitman coalescent given by Goldschmidt and Martin in [6]; then we follow the argument given in the same paper to establish that

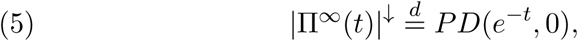

where *PD*(*α, θ*) is the (*α, θ*)−Poisson-Dirichlet distribution.

Briefly, the construction of the Bolthausen-Sznitman coalescent in terms of Random Recursive Trees proceeds as follows. We work on the set of recursive trees whose labeled nodes form a partition *π* of [*n*] := {1, …, *n*}, where the ordering of the nodes that confers the term “recursive” is given by ordering the blocks of *π* according to their least elements. A cutting-merge procedure is defined on the set of recursive trees of this form with a marked edge, this procedure consists of cutting the marked edge and merging all the labels in the subtree below with the node above, thus creating a new recursive tree whose labels form a new (coarser) partition of [*n*] (see Figure 2). With this operation in mind we consider a RRT with labels {1}, …, {*n*}, say *T*, to which we also attach independent standard exponential variables to each edge. Then, for each time *t* > 0 we retrieve the partition of [*n*] obtained by performing a cutting-merge procedure on all the edges of *T* whose exponential variable is less than *t*. This gives a stochastic process (Π^*n*^(*t*))_*t*≥0_ with values on the set of partitions of [*n*] that can be proven to be the *n*-Bolthausen-Sznitman coalescent.

**Figure 2:**
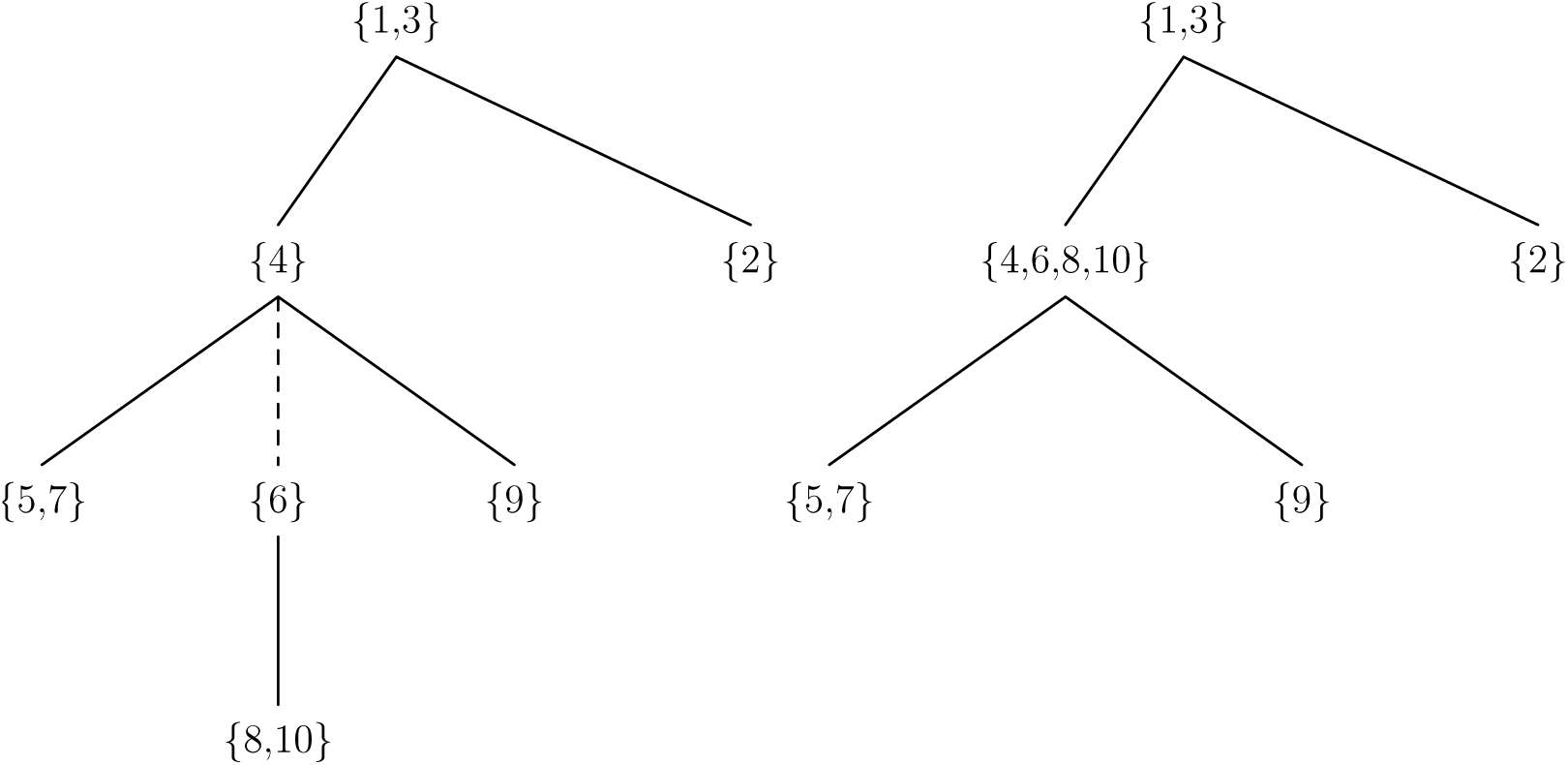
On the left, an example of a recursive tree whose labels constitute a partition of {1, …, 10}. On the right, the resulting recursive tree after a cutting-merge procedure performed on the marked edge (dashed line) of the first tree.

**Figure 3:**
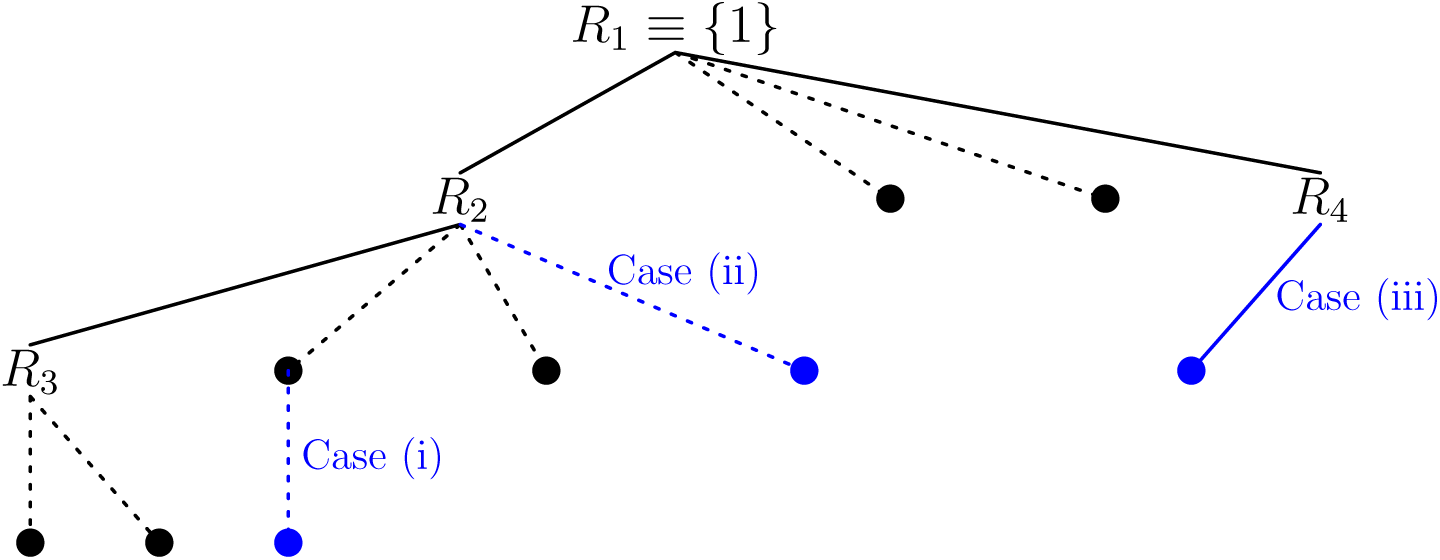
Schematic representation of passing from Π^*n*^(*t*) to Π^*n*+1^(*t*) for fixed *t*, by adding a new node (blue) to a RRT. Solid lines and dotted lines represent edges whose exponential variables are greater than *t* and less than or equal to *t*, respectively. In this case at time *t* there are four subtrees rooted at *R*_1_, *R*_2_, *R*_3_, and *R*_4_, which form the blocks that constitute Π^*n*^(*t*); these blocks are also the tables of a Chinese Restaurant Process. In case (i) the new node will be included in the block formed by *R*_2_ at time *t*, irrespective of whether its exponential edge is greater than *t* or not. In case (ii) the new node forms part of the block rooted at *R*_4_ because its exponential edge is less than *t*. Finally, in case (iii) the new node is a new root of a subtree that will form an additional block of Π^*n*+1^(*t*) (i.e. the new node opens a new table in the Chinese Restaurant Process).

**Figure 4:**
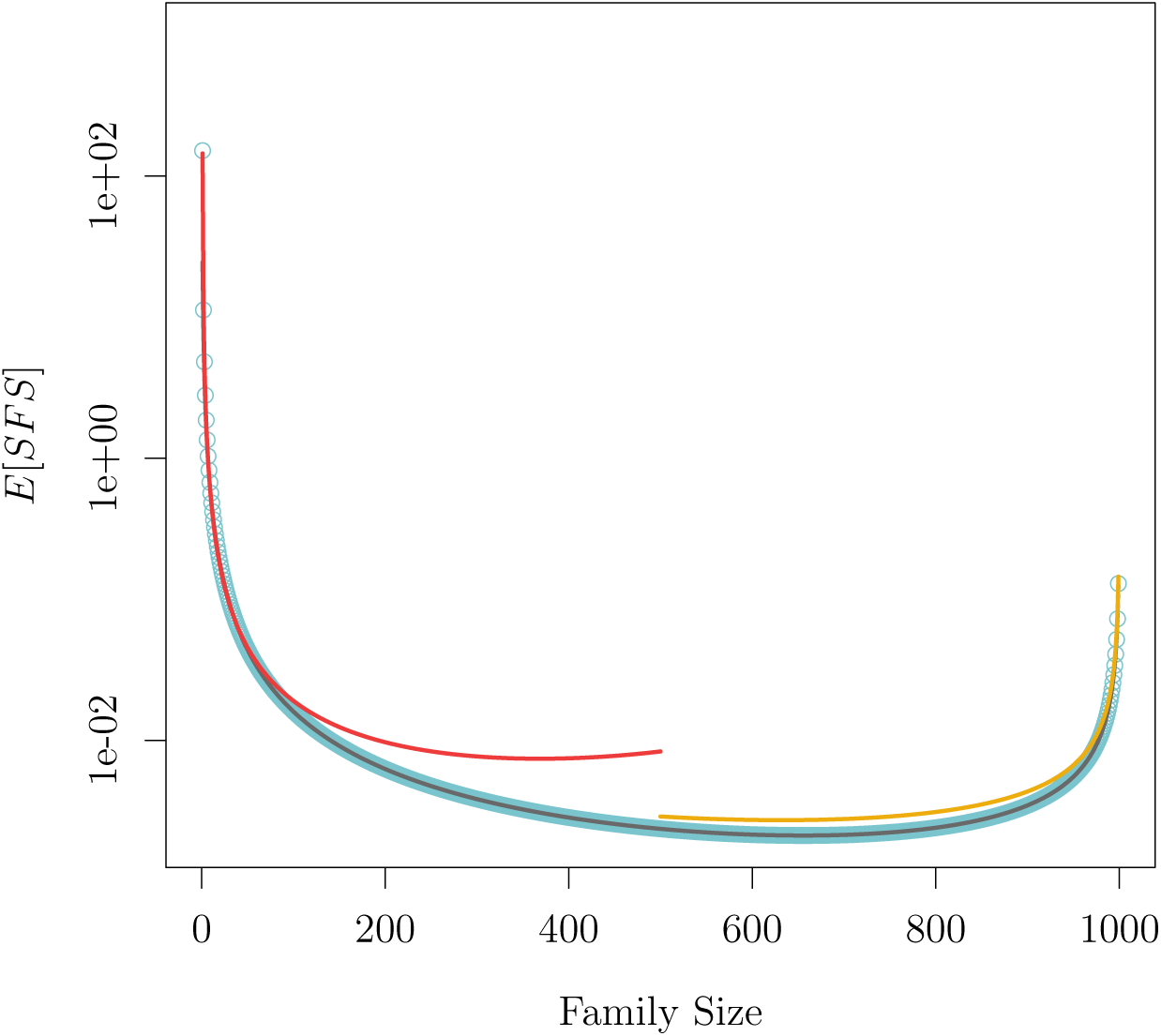
Exact and asymptotic approximations for 𝔼 [*SFS*] in a population of size 1000: The blue circles give the exact value as given in Corollary 3.3. The gray line is the asymptotic approximation as given in Theorem 3.4 (iii). Red (resp. yellow) line is given by Theorem 3.4 (ii) (resp. (iv)).

The fact that 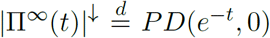 now follows readily. To see this, consider the construction of *T* where nodes arrive sequentially and each arriving node attaches to any of the previous nodes with equal probability. Considering also their exponential edges and having in mind the cutting-merge procedure we see that for any fixed time *t*, and assuming that *b* − 1 nodes have arrived and formed *k* blocks of sizes *s*_1_, …, *s*_*k*_ in Π^*b*−1^(*t*), the next arriving node, node {*b*}, will form a new block in Π^*b*^ (*t*) if and only if it attaches to any of the roots of the sub-trees of *T* that form the said *k* blocks and if, furthermore, its exponential edge is greater than *t*; this occurs with probability 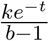. On the other hand, in order for *{b}* to join the *jth* block of size *s*_*j*_ it must either attach to the root of the sub-tree of *T* that builds this block and its exponential edge must be less than *t*, which happens with probability 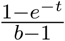, or it must attach to any other node of the said sub-tree, which happens with probability 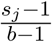; thus, the probability of attaching to the *jth* block is 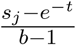. We recognize in these expressions the probabilities that define the Chinese Restaurant Process with parameters *α* = *e*^−*t*^ and *θ* = 0.

We now provide two straightforward applications of the RRT construction described above which nonetheless contain the essential intuitions underlying the forthcoming proofs.

### 2.1 Site Frequency Spectrum in the infinite coalescent

For the first application consider a subset *I* ⊂ (0, 1) and define (*C*_*I*_(*t*))_*t*≥0_ to be the process of the number of blocks in Π^∞^(*t*) with asymptotic frequencies in *I*. Then

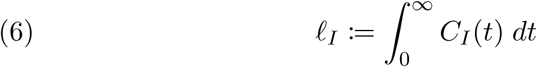

gives the total branch length of families with size frequencies in *I* in the infinite coalescent.

Our first theorem is a simple corollary of the equality in law (5).

#### Theorem 2.1.

For *I* ⊂ (0, 1), we have

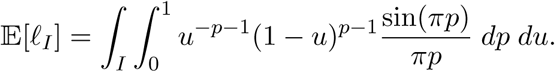

In particular, note that if in the infinite sites model with mutation rate *θ* we define *SFS*_*I*_ to be the number of mutations shared by a proportion *u* of individuals with *u* ranging in *I*, then by conditioning on *ℓ;*_*I*_ we get

#### Corollary 2.2.

For *I* ⊂ (0, 1), we have

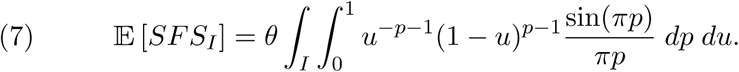

*Proof of* *Theorem* 2.1. Since

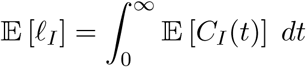

it only remains to compute 𝔼 [*C*_*I*_(*t*)] and simplify the expressions, but this is a straightforward consequence of Equation (6) in [14] which states that if *ϱ* = (*a*_1_, …) is *PD*(*α, θ*) distributed, and *f* : ℝ →ℝ is a function, then

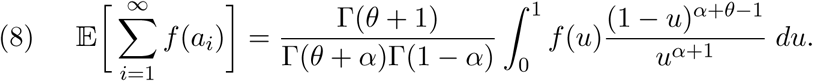

Taking *f* (*u*) = 𝟙 *I* (*u*)we get

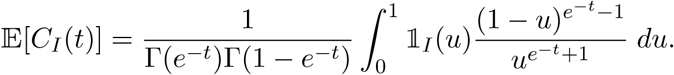

Using Euler’s reflection formula, making *p* = *e*^−*t*^ on the above expression and integrating on [0, ∞) we finish the proof. □

### 2.2 Time to the absorption

In this section we prove a useful lemma for the upcoming proofs, but a first consequence of this lemma gives the distribution function of the time to absorption, *A*_*n*_, in the *n*-coalescent, a result already proved in [12].

Here *Be* stands for the Beta function

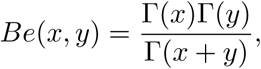

and Ψ for the digamma function

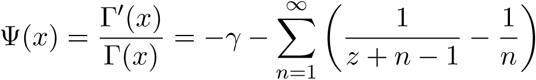

where *γ* stands for the Euler-Mascheroni constant.

#### Lemma 2.3.

Let *T* be a RRT on a set of *n* labels and with exponential edges. Define the two functionals *m*(*T*) and *M* (*T*) that give the minimum and the maximum of the exponential edges attached to the root of *T*. Then

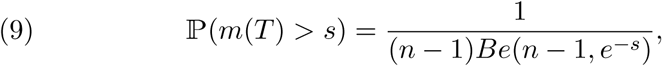

and

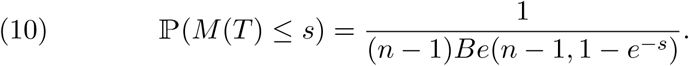

Also, for independent trees *T*_1_ and *T*_2_ of respective size *n*_1_ and *n*_2_, we have

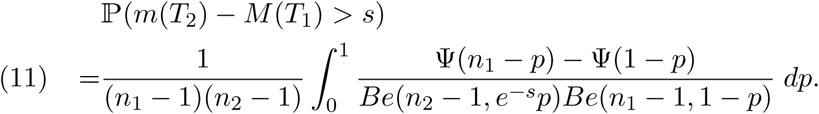

The proof of (10) follows the same lines as in [12] where the law of the time to absorption of the Bolthausen-Sznitman coalescent is derived, since this time is the maximum of the exponential edges attached to the root of a RRT. That is,

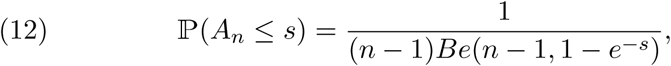

and, as *n* → ∞,

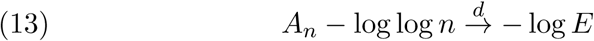

where *E* is a standard exponential random variable. The latter convergence in distribution was elegantly proved in [6] using a construction of random recursive trees in continuous time, whereas in this case it follows from Stirling’s approximation to the Gamma functions appearing in (12).

On the other hand, the equality (11) will be used in the computation of the distribution function of branch lengths with large family sizes presented in Section 4.

*Proof of Lemma* 2.3. Let *E*_2_, …, *E*_*n*_ be the exponential edges associated to the nodes of *T*. For the proof of (9) we consider the event {*m*(*T*) > *s*}. This event occurs when, in the recursive construction of *T* along with the exponential edges, the *i*th node (2 ≤ *i* ≤ *n*) does not attach to {1} whenever *E*_*i*_ < *s*; this happens with probability 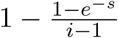. Thus, considering the *n* nodes, we obtain

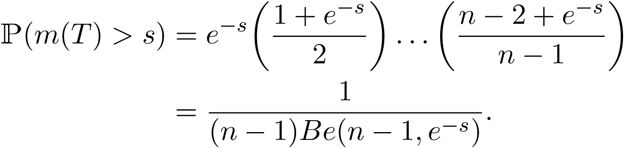

For (10) we instead build the tree such that the *i*th node does not attach to {1} whenever *E*_*i*_ > *s*; this happens with probability 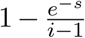. Thus we obtain

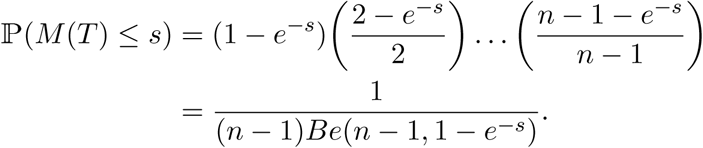

Finally we compute

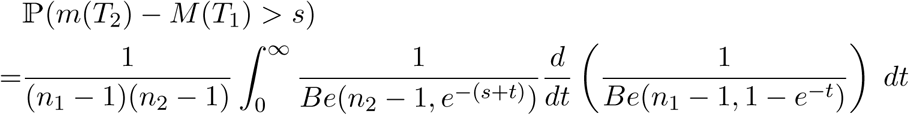

and by changing the variable *p* = *e*^−*x*^ we obtain (11). □

## 3 Moments of the Site Frequency Spectrum

By a simple adaptation of our previous notation for branch lengths in the infinite coalescent (*C*_*I*_ and *ℓ* _*I*_), in the finite case we also define for 1 ≤ *b* ≤ *n* − 1 the process (*C*_*n,b*_(*t*))_*t*≥0_ and the random variables (*ℓ* _*n,b*_), where *C*_*n,b*_(*t*) is the number of blocks of size *b* in Π^*n*^ (*t*), and

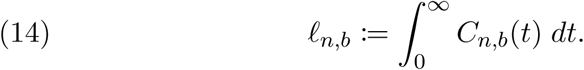

We now provide explicit expressions for 𝔼 [*ℓ*_*n,b*_] and 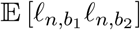; for this we define the functions

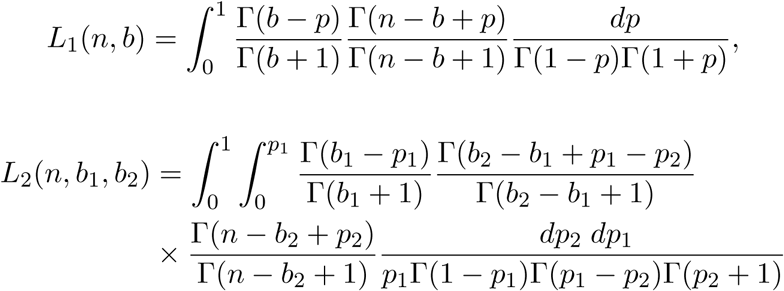

and

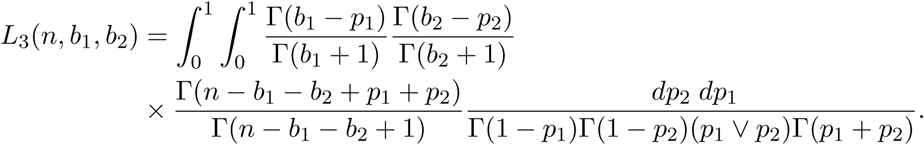

### Theorem 3.1.

For any pair of integers *n, b* such that 1 ≤ *b* ≤ *n* − 1, we have

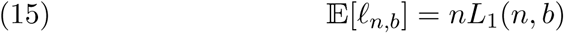

Also, for any triple of integers *n, b*_1_, *b*_2_, with 1 ≤ *b*_1_ ≤ *b*_2_ ≤ *n* − 1, we have

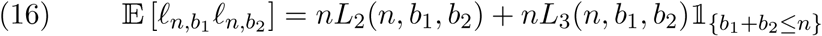

As before, we may define *SFS*_*n,b*_ as the number of mutations shared by *b* individuals in the *n*-coalescent. By conditioning on the value of the associated branch lengths we get

### Corollary 3.2.

For 1 ≤ *b* ≤ *n* − 1,

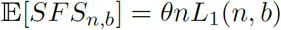

and, for 1 ≤ *b*_1_ ≤ *b*_2_ ≤ *n* − 1, we have,

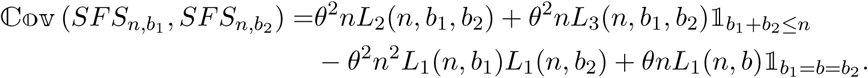

We also characterize the asymptotic behavior of the functions *L*_1_, *L*_2_ and *L*_3_ as *n* → ∞, which in turn give asymptotic approximations for the first and second moments of the branch lengths and of *SFS*. For this we recall the function *f*_1_ defined in (2) and also define for 0 < *u*_1_ < *u*_2_ < 1,

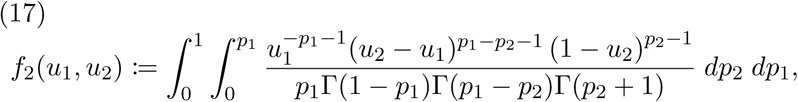

and, for *u*_1_, *u*_2_ > 0, *u*_1_ + *u*_2_ < 1,

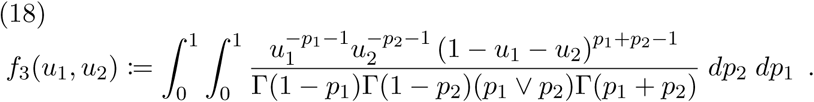

### Lemma 3.3.

We have as *n* → ∞,

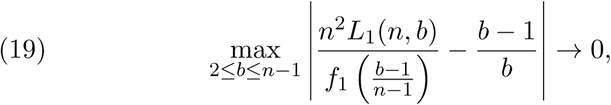

whereas for *b* = 1,

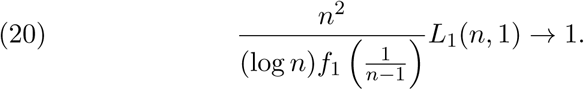

Similarly

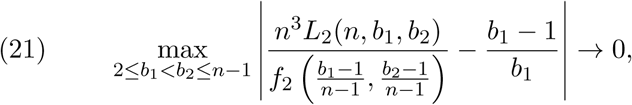

and if also *b*_1_ ∨ (*n* − *b*_2_) → ∞ then

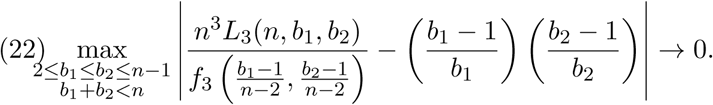

### Remark.

The above lemma does not cover the cases *b*_1_ = 1 or *b*_1_ = *b*_2_ for *L*_2_, nor the cases *b*_1_ = 1, *b*_2_ = 1, *n* = *b*_1_ +*b*_2_ or *b*_1_ ∨(*n*−*b*_2_) ↛ ∞ for *L*_3_. However, using the same techniques we also obtain asymptotics in these cases which are used in Theorem 3.5 below.

The proof of the above lemma also gives asymptotic expressions for the functions *f*_1_, *f*_2_ and *f*_3_, leading to straightforward asymptotics for the expectation and covariance of *SFS*. The complete picture for the first moment is given in the next result.

### Theorem 3.4.

As *n* goes to infinity,

i. The expected number of external mutations (*b* = 1) has the following asymptotics

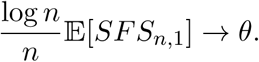
ii. If *b* ≥ 2 and 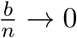, then

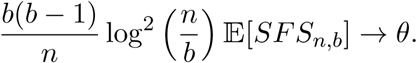
iii. If 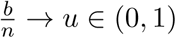, then

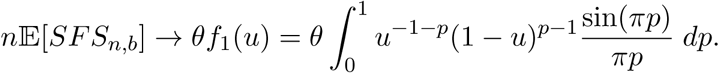
iv. If 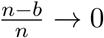, then

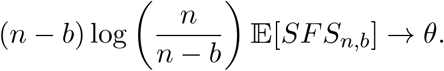
v. Let *I* = (*x, y*) with 0 < *x* < *y* < 1 and define

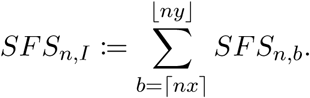

Then

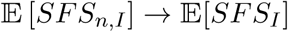

as it is defined in (7).

Case (i) and case (ii) for fixed *b* also follow from Theorem 4 in [2]. Cases (ii) and (iv) give an update to the approximation of the SFS for small and large families made in [13].

In the same spirit and using the same techniques we now provide the complete picture for the second moments. In what follows we use the notation *f* (*n*) ∼ *g*(*n*) to denote that

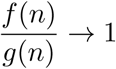

as *n* → ∞.

### Theorem 3.5.

The covariance function has the following asymptotics as *n* goes to infinity, in each of the following cases:

| $b_1$ | $b_2 - b_1$ | $n - b_2$ | $\mathbb{C}\mathbb{O}\mathbb{V}(SFS_{n,b_1}, SFS_{n,b_2})$ |
| --- | --- | --- | --- |
| $> 1$ | $> 0$ | $\sim n$ | $\frac{\theta^2}{b_1(b_1-1)b_2(b_2-1)} \mathcal{O}\left(\frac{n^2}{\log^5 n}\right)$ |
| $\sim n$ | $> 0$ | $> 0$ | $\frac{\theta^2}{(b_2-b_1)(n-b_1)} \frac{1}{\log^2 n}$ |
| $\sim n$ | $0$ | $> 0$ | $\frac{\theta^2 + \theta}{n-b_2} \frac{1}{\log n}$ |
| $> 1$ | $\sim n$ | $= b_1$ | $\theta^2 \mathcal{O}\left(\frac{n}{\log^4 n}\right)$ |
| $> 1$ | $\sim n$ | $= b_1 + \text{const}^+$ | $\theta^2 L_1(n - b_2, b_1) \frac{n}{\log n}$ |
| $1$ | $0$ | $\sim n$ | $\theta^2 \mathcal{O}\left(\frac{n^2}{\log^3 n}\right)$ |
| $1$ | $> 0$ | $\sim n$ | $\theta^2 \mathcal{O}\left(\frac{n^2}{\log^4 n}\right)$ |
| $1$ | $\sim nu$ | $\sim n(1-u)$ | $\theta^2 \mathcal{O}\left(\frac{1}{\log^2 n}\right)$ |
| $1$ | $\sim n$ | $> 1$ | $\theta^2 \mathcal{O}\left(\frac{n}{\log^3 n}\right)$ |
| $1$ | $\sim n$ | $1$ | $\theta^2 \mathcal{O}\left(\frac{n}{\log^3 n}\right)$ |
| $> 1$ | $0$ | $\sim n$ | $\theta^2 \mathcal{O}\left(\frac{n^2}{\log^5 n}\right)$ |
| $\sim nu$ | $> 0$ | $\sim n(1-u)$ | $\frac{\theta^2}{(1-u)(b_2-b_1)} \frac{1}{n \log^2 n}$ |
| $\sim nu$ | $0$ | $\sim n(1-u)$ | $\frac{\theta f_1(u)}{n}$ |
| $> 1$ | $\sim nu$ | $\sim n(1-u)$ | $\theta^2 \mathcal{O}\left(\frac{1}{\log^3 n}\right)$ |
| $\sim nu$ | $\sim n(1-u)$ | $> 0$ | $-\frac{\theta^2 f_1(u)}{n-b_2} \frac{1}{\log n}$ |
| $\sim nu_1$ | $\sim nu_2$ | $\sim n(1-u_1-u_2)$ | $\frac{\theta^2 (f_2(u_1, u_1+u_2) + f_3(u_1, u_1+u_2) \mathbb{1}_{2u_1+u_2 \leq 1} - f_1(u_1)f_1(u_1+u_2))}{n^2}$ |
| $\sim nu$ | $\sim n(1-2u)$ | $= b_1$ | $\frac{\theta^2 \int_0^\infty \int_0^\infty \frac{e^{-y_1} e^{-y_2}}{y_1 \vee y_2} dy_1 dy_2}{u(1-u)} \frac{1}{n \log n}$ |
| $\sim nu$ | $\sim n(1-2u)$ | $= b_1 + \text{const}^+$ | $\frac{\theta^2 \int_0^\infty \int_0^\infty \frac{e^{-y_1} e^{-y_2} (y_1+y_2)}{y_1 \vee y_2} dy_1 dy_2}{u(1-u)(n-b_2-b_1)} \frac{1}{n \log^2 n}$ |

Also for *I, Î* ⊂ (0, 1), and *SFS*_*n,I*_, *SFS* _*n, Î*_ as defined in Theorem 3.4 (V), we have

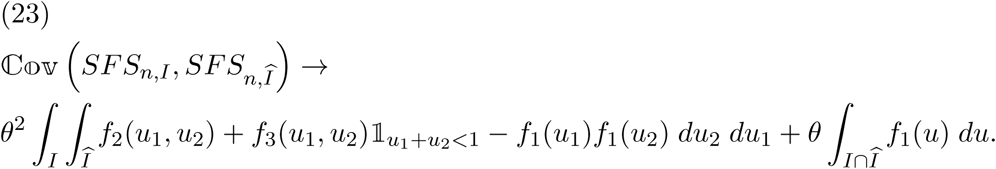

These approximations follow from the asymptotics for *L*_1_, *L*_2_, and *L*_3_ substituted in the covariance formula given in Corollary 3.3. For the sake of simplicity we do not provide the explicit computations. We only treat the case where the expected value 𝔼[*SFS*_*n,b*_] diverges, then an application of Chebyshev’s inequality allows us to prove the following weak law of large numbers with *L*^2^-convergence, which generalizes and strengthens results on the Bolthausen-Sznitman coalescent derived in [2].

### Corollary 3.6.

Suppose that *b*/*n* → 0 in such a way that 𝔼[*SFS*_*n,b*_] →∞, or equivalently that 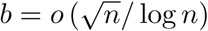. Then we have the following *L*^2^-convergence:

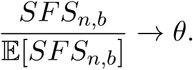

In view of Theorem 3.4 this means that for *b* = 1

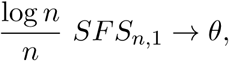

and for *b* ≥ 2, 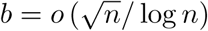

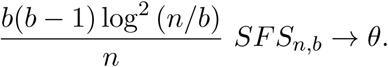

## 4 Distribution of the Family-Sized Branch Lengths

In this section we discuss the particular case of *ℓ*_*n,b*_ when *b* > *n/*2. In this case we are able to provide an explicit formula for the distribution function of the length of the coalescent of order *b*. This leads to convergence in law results, but also to the law of *SFS*_*n,b*_. Observe that in this case, for all *t* ≥ 0, *C*_*n,b*_(*t*) ∈ {0, 1} and *ℓ*_*n,b*_ is just the time during which the block of size *b* survives before coalescing with other blocks (if it ever exists, otherwise obviously *ℓ*_*n,b*_ = 0). We first find an expression for the distribution function of *ℓ*_*n,b*_.

### Theorem 4.1.

Suppose that *n/*2 < *b* < *n*. For any *s* ≥ 0,

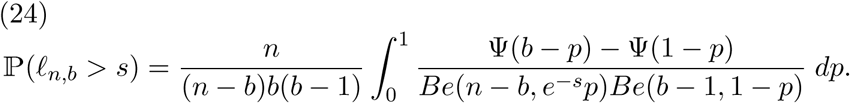

From the derived distribution of *ℓ*_*n,b*_ in Theorem 4.1 we obtain that, conditioned on *ℓ*_*n,b*_ > 0, the variable (log *n*) *ℓ*_*n,b*_ has a limiting distribution.

### Corollary 4.2.

Suppose that *b/n* → *u* ∈ [1*/*2, 1) as *n* → ∞, then letting *α* = log(1 − *u*) − log *u*, we have

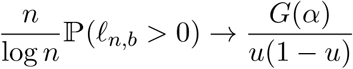

where

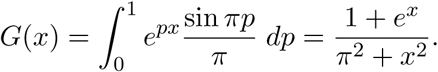

Furthermore,

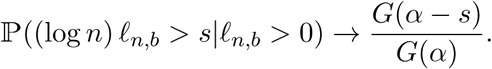

We now give the joint distribution of the branch lengths for large families, i.e. the joint distribution of the vector (*ℓ*_*n,b*_)*b*>*n/*2. For this we introduce the following events: for any collection of integers **b** = (*b*_1_, …, *b*_*m*_) such that *n/*2 < *b*_1_ < *b*_2_ < … < *b*_*m*_ < *n*, and any collection of nonnegative numbers **s** = (*s*_1_, …, *s*_*m*_), define the event

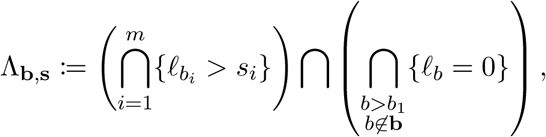

that is, the event that a block of size *b*_1_ exists for a time larger than *s*_1_, that this block then merges with some other blocks of total size exactly *b*_2_ − *b*_1_, that this new block exists for a time larger than *s*_2_, and so on, until the last merge of the growing block occurs with the remaining blocks of total size exactly *n* − *b*_*m*_.

### Theorem 4.3.

For **b** = (*b*_1_, …, *b*_*m*_) and **s** = (*s*_1_, …, *s*_*m*_) as above, we have

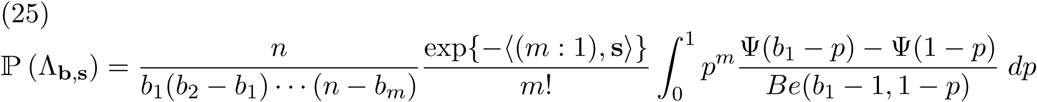

and

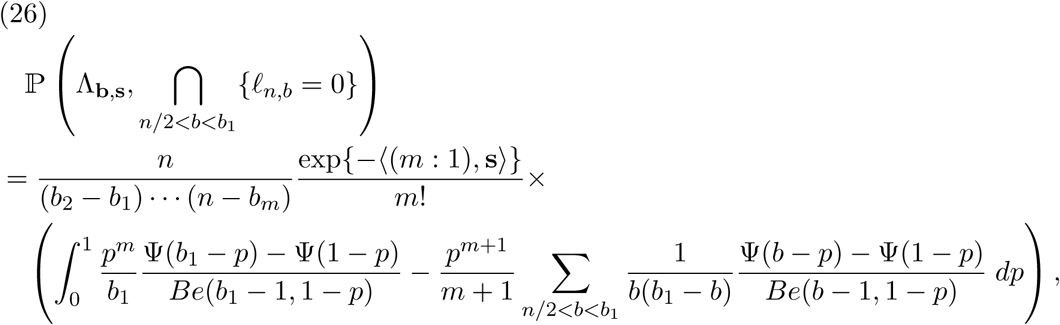

where

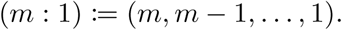

and ⟨·, ·⟩ is the usual inner product in Euclidean space.

By conditioning on (*ℓ*_*n,b*_)_*b*>*n/*2_ and using equation (26) one can obtain a sampling formula for the vector (*SFS*_*n,b*_)_*b*>*n/*2_, although the computations are rather convoluted and we do not present them here.

## 5 The approximations

Here we derive the approximations given above in the Introduction. From Stirling’s approximation we have the well-known formula Γ(*m* + *c*)*/*Γ(*m*) ≈*m*^*c*^. Its application requires some care, since we shall apply this approximation also for small values of *m* down to *m* = 1. It is known and easily confirmed by computer that the approximation is particularly accurate within the range 0 ≤ *c* ≤ 1. Thus we use for *p* ∈ (0, 1) and *b* ≥ 2 the approximations

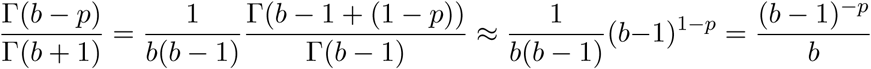

and

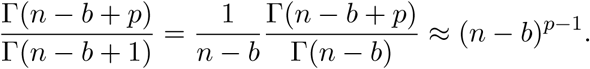

Also by Euler’s reflection formula Γ(1 − *p*)Γ(1 + *p*) = *πp/* sin(*πp*). Inserting these formulas into the expression (15) for the expected SFS we obtain

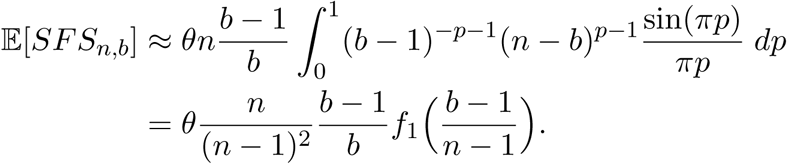

It turns out that this approximation overestimates the expected SFS, which can be somewhat counterbalanced by replacing the scaling factor *n/*(*n* − 1)^2^ by 1*/*(*n* − 1). This yields our first approximation (1). For the second approximation (3) we apply the expansion

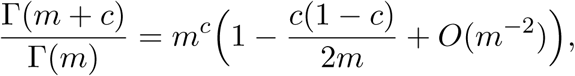

see [18]. Again this approximation is particularly accurate for 0 ≤ *c* ≤ 1 leading for *p* ∈ (0, 1) and *b* ≥ 2 to

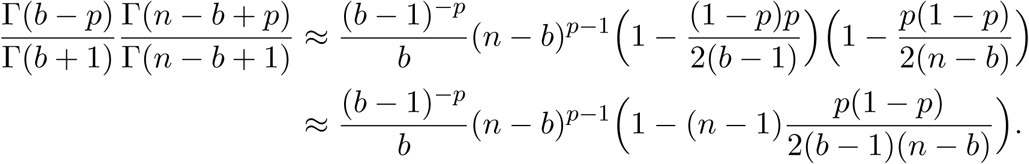

Using this approximation in the expression for the expected SFS we get for *b* ≥ 2

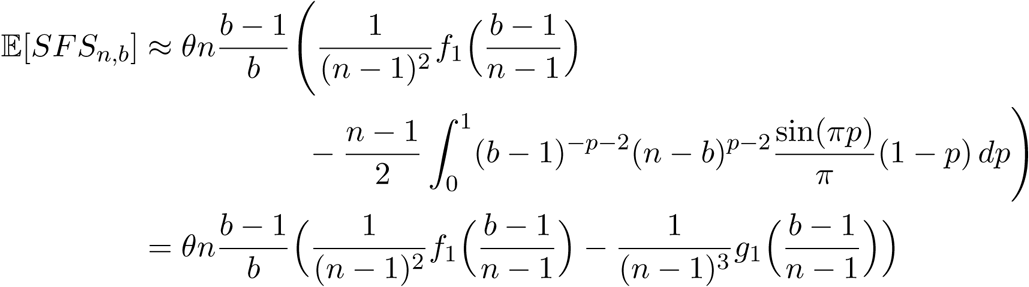

with the function *g*_1_ as defined in (4). This integral can be evaluated by elementary means yielding formula (3).

## 6 Proofs of Section 3

As in the infinite coalescent case, the proof of Theorem 3.1 begins with the definition (14) and by noting that

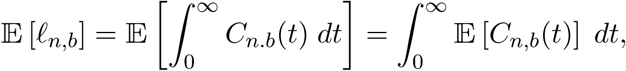

and similarly

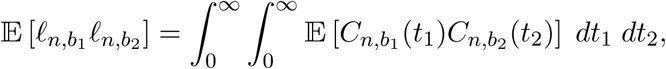

so it only remains to compute 𝔼 [*C*_*n,b*_(*t*)] and 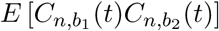 in each case and simplify the expressions.

### Proof of Theorem 3.1

*(first moment).* Let ℬ be the collection of all possible blocks of size *b* in a partition of [*n*]. Then

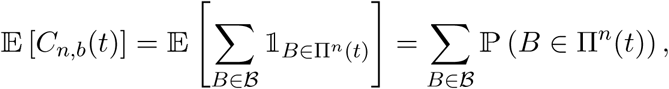

and by exchangeability of Π^*n*^(*t*),

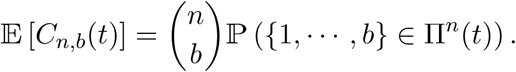

Thus, using (8), the fact that 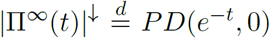, and writing Π^*n*^ as 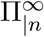, we obtain

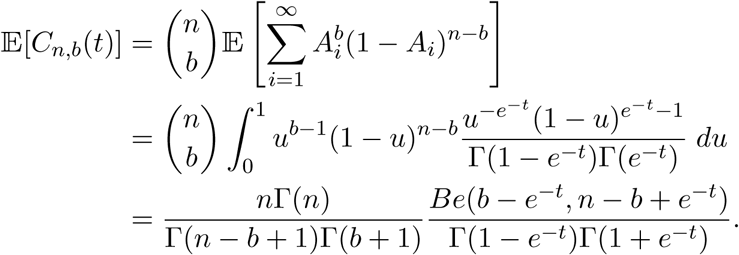

Finally, by changing the variable *p* = *e*^−*t*^, we obtain (15). □

Now we use the random tree construction of the *n*-Bolthausen-Sznitman coalescent in order to compute the second moments of 𝓁_*n,b*_.

### Proof of Theorem 3.1

*(second moments).* Let 1 ≤ *b*_1_ ≤ *b*_2_ ≤ *n* − 1, and ℬ _1_, ℬ_2_ be the collection of all possible blocks of sizes *b*_1_ and *b*_2_ respectively in a partition of [*n*]. Then

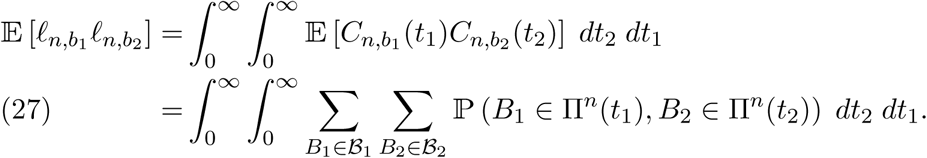

We now compute ℙ (*B*_1_ ∈ Π^*n*^(*t*_1_), *B*_2_ ∈ Π^*n*^(*t*_2_)) by cases.

i) Suppose that *B*_1_ ∩ *B*_2_ = ∅. By exchangeability we have

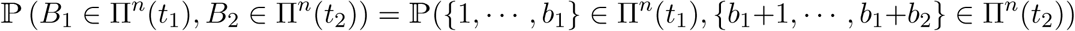

where this probability is of course 0 if *b*_1_ + *b*_2_ > *n*. Now suppose that *t*_1_ ≤ *t*_2_. In terms of the RRT construction of the Bolthausen-Sznitman coalescent, the event

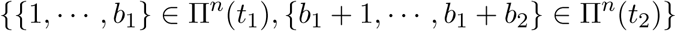

is characterized by a RRT with exponential edges, say *E*_2_, …, *E*_*n*_, constructed as follows: for *i* ∈ {1, …, *b*_1_ − 1} the node {*i* + 1} along with *E*_*i*+1_ arrive to the tree but with the imposed restriction that it may not attach to {1} and have *E*_*i*+1_ > *t*_1_ at the same time, which occurs with probability 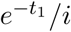; this ensures that {*i* + 1} coalesces with {1} before time *t*_1_ for all *i* < *b*_1_, thus creating the block {1, …, *b*_1_} up to time *t*_1_. After {1}, …, {*b*_1_} have arrived, the node {*b*_1_ + 1} must attach to {1} and 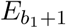 must be greater than *t*_2_, which occurs with probability 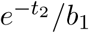; the node {*b*_1_ + 1} will be the root of a sub-tree formed with the nodes {*b*_1_ +2}, …, {*b*_1_ +*b*_2_} which will build the block {*b*_1_ + 1, …, *b*_1_ + *b*_2_} at time *t*_2_. Thus, for each *i* ∈ {1, …, *b*_2_ − 1} the node {*b*_1_ +*i*+1} must arrive and attach to any of {*b*_1_ +1}, …, {*b*_1_ +*i*}, which occurs with probability 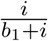, and, furthermore, conditional on this event, it may not attach to {*b*_1_ + 1} and have 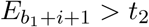 at the same time, which occurs with probability 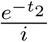. Finally, if *n* − *b*_1_ − *b*_2_ > 0, for *i* ∈ {0, …, *n* − *b*_1_ − *b*_2_ − 1} the node {*b*_1_ + *b*_2_ + *i* + 1} must either attach to any of {*b*_1_ + *b*_2_ + *j*}, 1 ≤ *j* ≤ *i*, or attach to {1} or {*b*_1_ + 1} and have 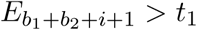 or 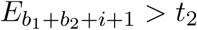 respectively; this occurs with probability 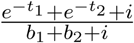. Putting all together we obtain

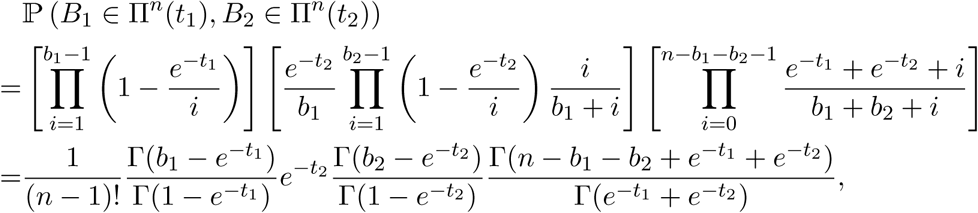

where the last product is set to 1 if *n* − *b*_2_ − *b*_1_ = 0. On the other hand, if *t*_2_ < *t*_1_, by exchangeability we may instead compute

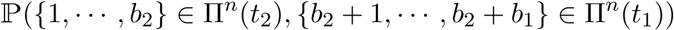

obtaining

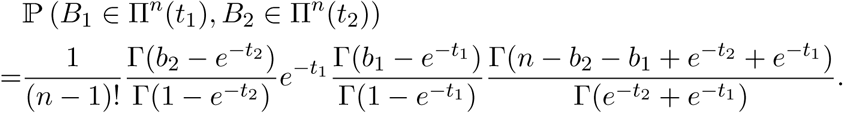

ii) Suppose that *B*_1_ ⊂ *B*_2_. Of course if *t*_1_ > *t*_2_ we have ℙ (*B*_1_ ∈ Π^*n*^(*t*_1_), *B*_2_ ∈ Π^*n*^(*t*_2_)) = 0 whenever *B*_1_ is strictly contained in *B*_2_. Assuming that *t*_1_ ≤ *t*_2_ and using the same rationale as before we obtain

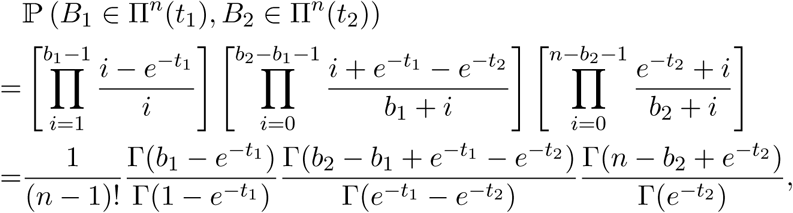

where the product in the middle is set to 1 if *B*_1_ = *B*_2_.

iii) If *B*_1_∩*B*_2_ ≠ ∅ and *B*_1_ ⊄ *B*_2_, we clearly have ℙ (*B*_1_ ∈ Π^*n*^(*t*_1_), *B*_2_ ∈ Π^*n*^(*t*_2_)) = 0.

From the previous computations, and summing over the corresponding cases, we see that if *b*_1_ + *b*_2_ ≤ *n* then, changing the variable *p* = *e*^−*t*^, the integral in (27) is given by

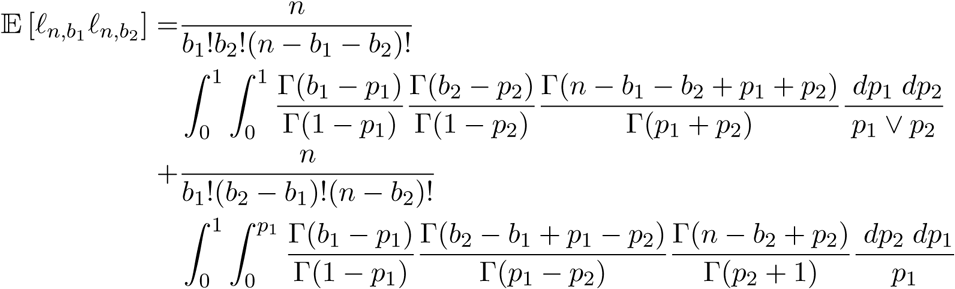

whereas if *b*_1_ + *b*_2_ > *n* the first summand in the above expression is set to zero. Rearranging terms we obtain (16). □

*Proof of Lemma* 3.3 *(asymptotics for L*_1_). Again, we have from Stirling’s formula that Γ(*m* + *c*)*/*Γ(*m* + *d*) = *m*^*c*−*d*^(1 + 𝒪 (1*/m*)) for any real numbers *c* and *d*, where the 𝒪 (1*/m*) term holds uniformly for 0 ≤ *c, d* ≤ 1. Letting *m* = *b* − 1 and *n* − *b* leads to the following equality:

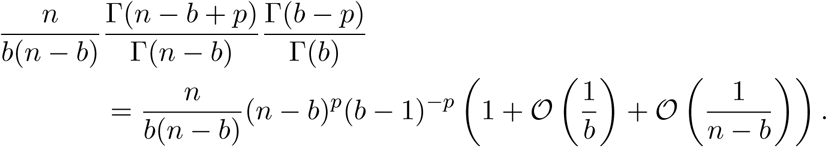

Thus, using Euler’s reflection formula to write Γ(1 − *p*)Γ(1 + *p*) as *πp/* sin (*πp*) in the definition of *L*_1_, we get

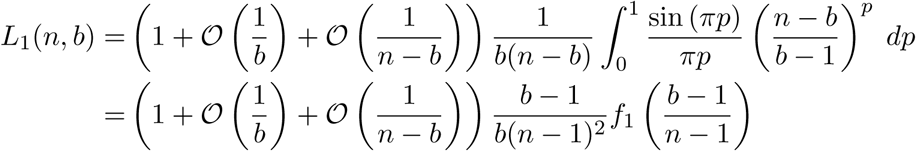

Thus, for every *ϵ* > 0 there is a *b*_0_ ∈ ℕ such that for large enough *n* ∈ ℕ we have

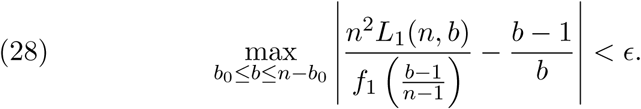

It remains to study the approximation as *n* → ∞ in the cases where *n* − *b* or *b* remain constant. In the first case, when *n* − *b* = *c*, we have *b* → ∞ as *n* → ∞ and, by Stirling’s approximation and dominated convergence and substituting *p* = *y/* log *b* on the one hand

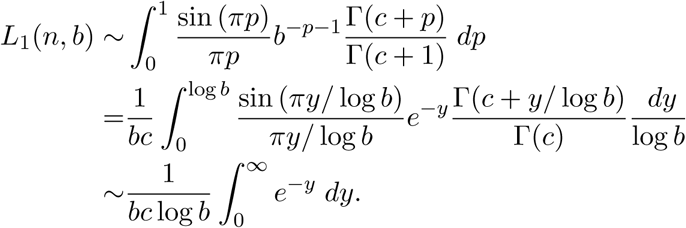

and on the other hand because of *b* → ∞

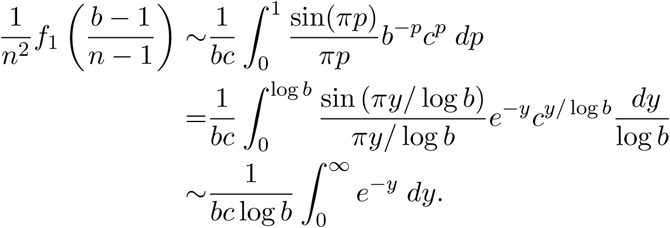

Thus *L*_1_(*n, b*) ∼ *n*^−2^*f*_1_((*b* − 1)*/*(*n* − 1)) which extends (28) for *b* > *n* − *b*_0_.

Similarly for the second case, if *b* ≥ 2 is fixed, we have *n* − *b* → ∞ as *n* → ∞. Thus, with 1 − *p* = *y/* log *n*

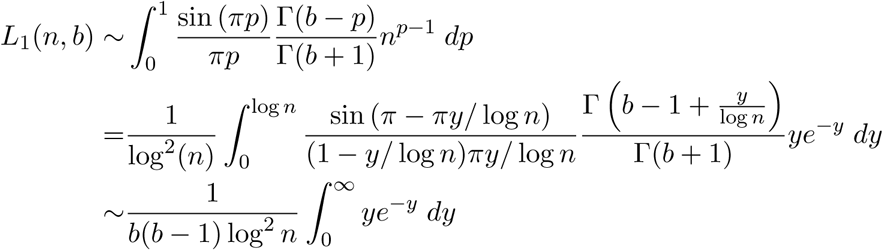

and

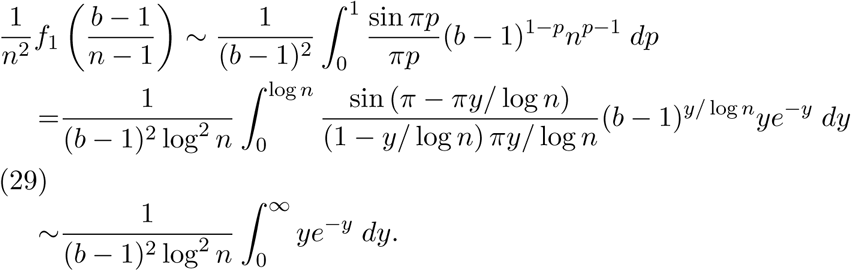

Thus *L*_1_(*n, b*) ∼ (*b* − 1)*n*^−2^*f*_1_((*b* − 1)*/*(*n* − 1))*/b*, which extends (28) for *b* < *b*_0_. This extends (28) for *b* < *b*_0_. Thus we proved (19).

For the proof of (20), we substitute *b* by 1 and perform similar computations:

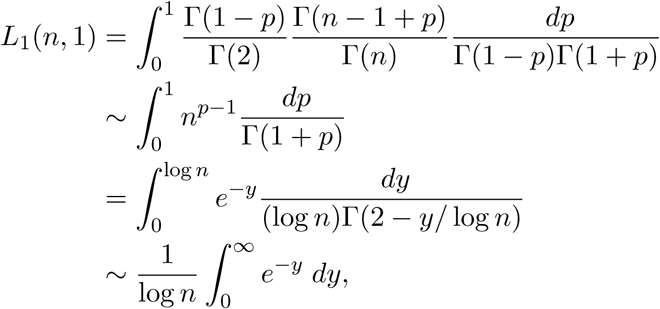

and from (29) with choosing *b* = 2

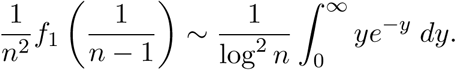

This proves (20). □

*Proof of Lemma* 3.3 *(asymptotics for L*_2_ *and L*_3_). The arguments here are similar to the arguments in the proof of the asymptotics for *L*_1_, but we avoid repeating similar and tedious computations. We only layout the first steps of the proof. By Stirling’s approximation applied to the integrands appearing in *L*_2_ and *L*_3_, we obtain, for *b*_2_ − *b*_1_ > 0,

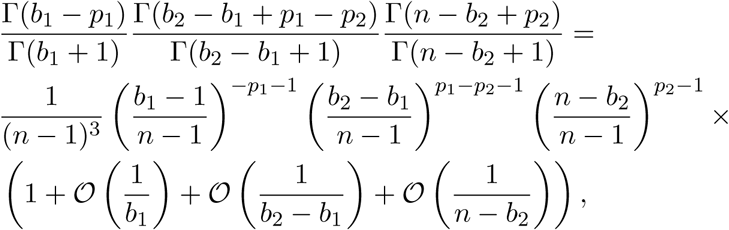

and, for *n* − *b*_2_ − *b*_1_ > 0,

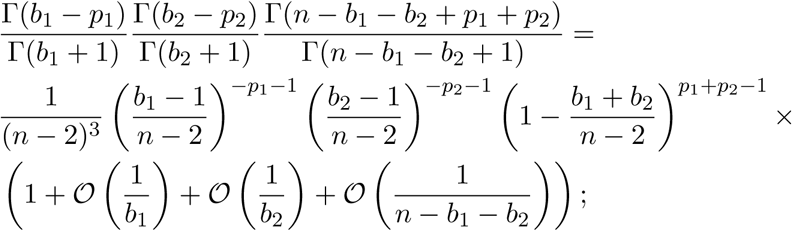

thus

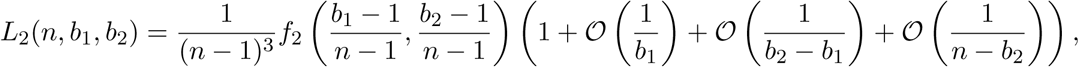

and

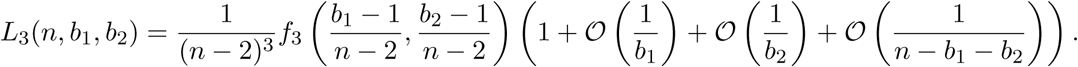

Similar to the analysis in the proof of (19), to obtain (21) it remains to study the cases where at least one of *b*_1_, *b*_2_ − *b*_1_, or *n* − *b*_2_ remains constant, whereas for (22) the cases of interest are where one of *b*_1_, *b*_2_, or *n* − *b*_2_ − *b*_1_ remain constant. □

*Proof of Theorem 3.4.* We first derive the asymptotic behavior of the function *f*_1_. We have

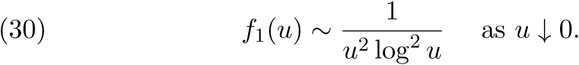

For the proof note that for *u* < 1*/*2 we have (1 − *u*)^*p*−1^ ≤ 2. Therefore dominated convergence implies for *u* ↓ 0

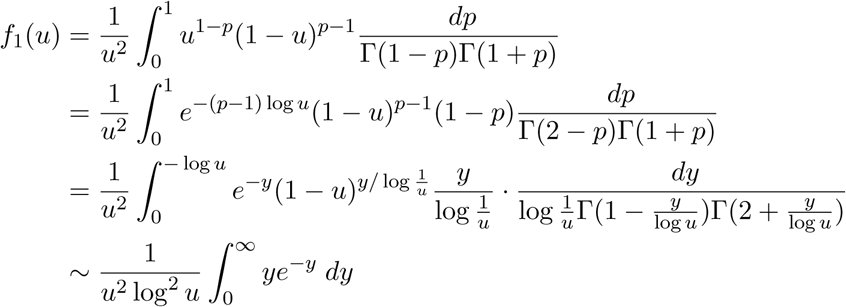

implying (30). Also

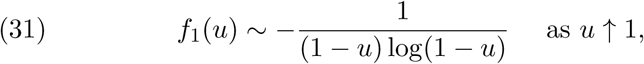

which we obtain again by means of dominated convergence in the limit *u* ↑ 1 as follows:

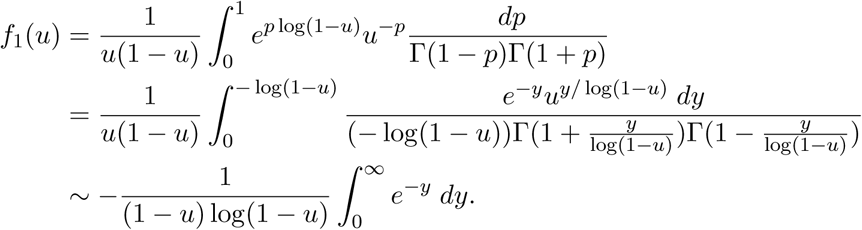

These asymptotics together with Lemma 3.3 imply our claims. Without loss of generality let *θ* = 1. From (20) we obtain

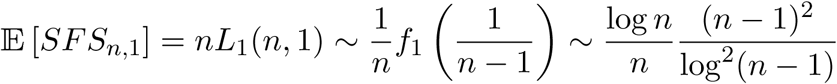

which yields claim (i).

Similary from (19) we get for *b* ≥ 2 and *b/n* → 0

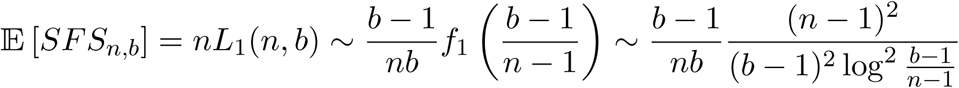

which in view of *b/n* → 0 yields assertion (ii).

Claim (iii) is an immediate consequence of formula (19), since here we have (*b* − 1)*/b* → 1.

Next under the condition (*n* − *b*)*/n* → 0 we get from (19) and (31)

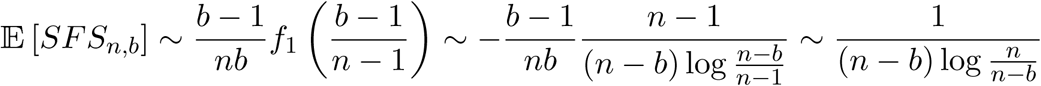

which confirms assertion (iv).

Finally, we have from (19)

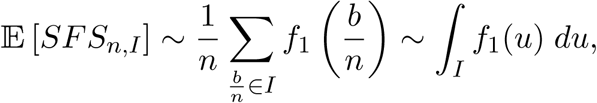

which is claim (v). This finishes the proof. □

### Proof of Theorem 3.5.

The approximations follow from the asymptotics for *L*_1_, *L*_2_, and *L*_3_ substituted in the covariance formula given in Corollary 3.3. □

*Proof of Corollary 3.3.* We have to prove that

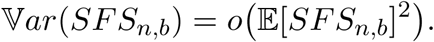

From the monotonicity properties of the gamma function we have for 1 ≤ *b* ≤ *n* – 1

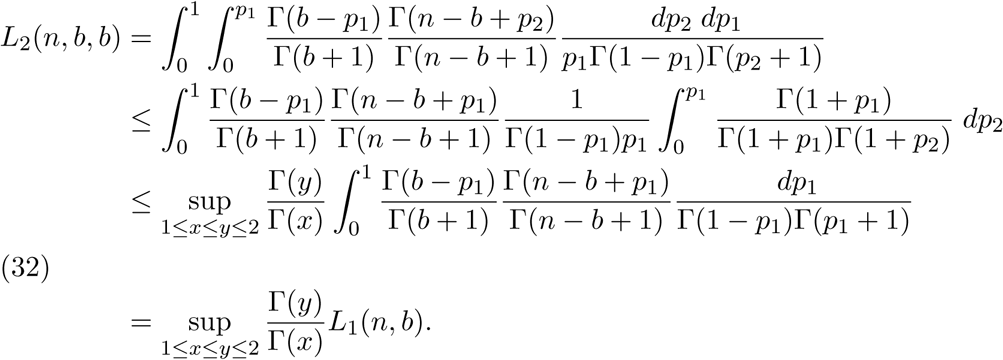

Concerning *L*_3_(*n, b, b*) we have for *b* = *o*(*n*) by Stirling’s approximation uniformly in 0 ≤ *p*_1_, *p*_2_ ≤ 1

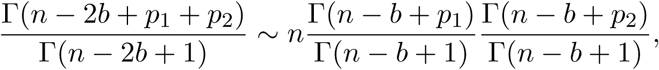

hence, with 1 < *η* < 2

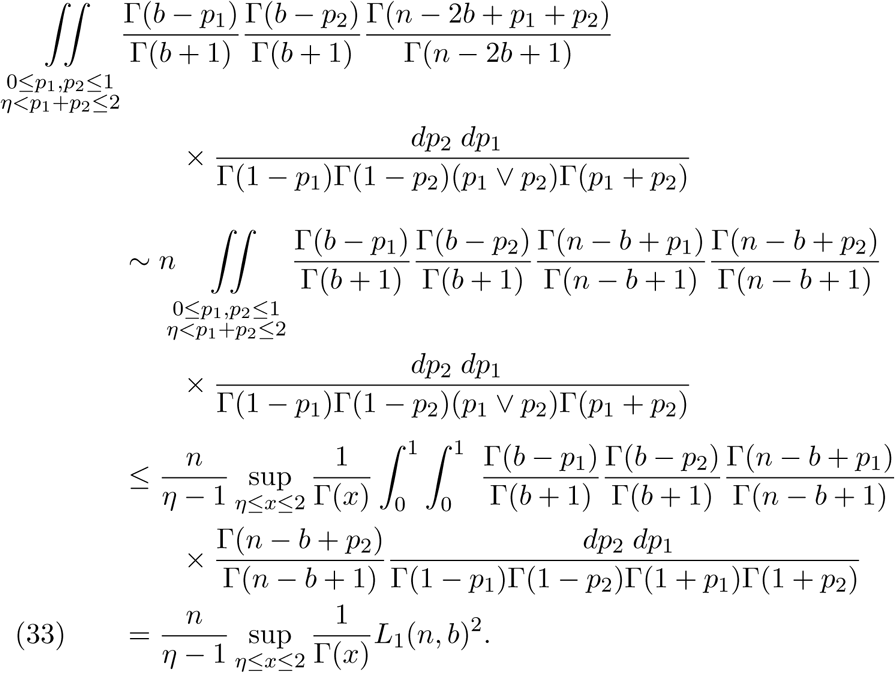

Also, by another application of Stirling’s approximation and for *b* = *o*(*n*)

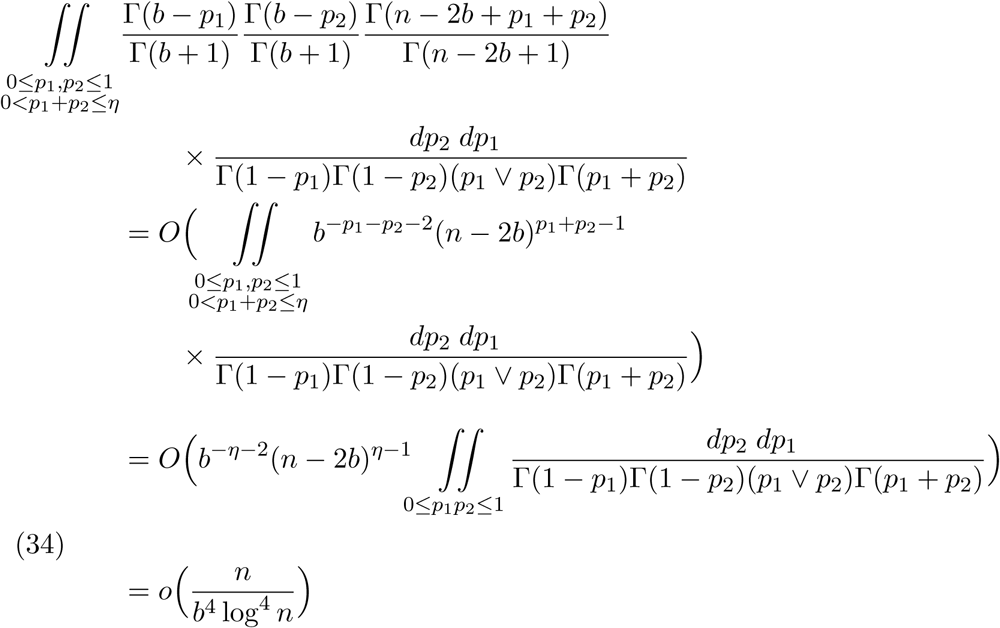

Combining (33) and (34) with Theorem 3.4 (i) and (ii) and letting *η* → 2 we obtain

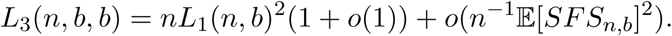

Using this estimate together with (32) and with Theorem 3.1, Corollary 3.3 yields

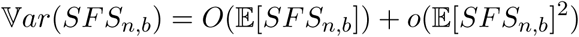

Because of our assumption 𝔼[*SFS*_*n,b*_] → ∞ our claim is proved. □

## 7 Proofs of Section 4

### Proof of Theorem 4.1.

Note that since *b* > *n/*2, and by the exchangeability of Π^*n*^, we have:

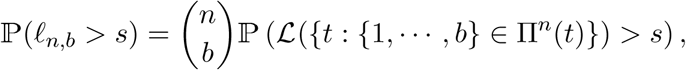

where ℒ is the Lebesgue measure, and ℒ({*t* : {1, …, *b*} ∈ Π^*n*^(*t*)}) gives the time that the block {1, …, *b*} exists in the Bolthausen-Sznitman coalescent starting with *n* individuals.

We now describe the event {ℒ({*t* : {1, …, *b*}} ∈ Π^*n*^(*t*)) > *s*} in terms of the RRT construction of the Bolthausen-Sznitman coalescent. Let 𝒢 be the event that the nodes {1}, {2}, …, {*b*} and {1}, {*b* + 1}, …, {*n*} form two sub-trees, say *T*_1_ and *T*_2_ rooted at {1}; i.e.

𝒢 :={*T* : {*j*} does not attach to {*i*}, for all 2 ≤ *i* ≤ *b* and *b* < *j* ≤ *n*}.

Then

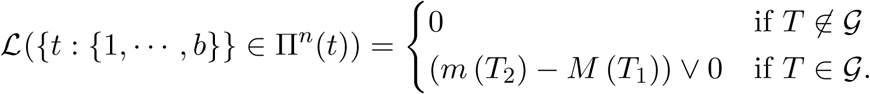

Indeed, observe that by the cutting-merge procedure *T* ∉ 𝒢 if and only if any block of Π^*n*^ that contains all of {1, …, *b*} also contains some *j* ∈ {*b* + 1, …, *n*}. On the other hand, on the event {*T* ∈ 𝒢}, the random variable *M* (*T*_1_) is just the time at which the block {1, …, *b*} appears in Π^*n*^, while *m*(*T*_2_) is the time at which it coalesces with some other block in *T*_2_. Furthermore, observe that conditioned on {*T* ∈ 𝒢}, *T*_1_ and *T*_2_ are two independent RRTs of sizes *b* and *n* − *b* + 1 respectively. Thus, by Lemma 2.3 we have

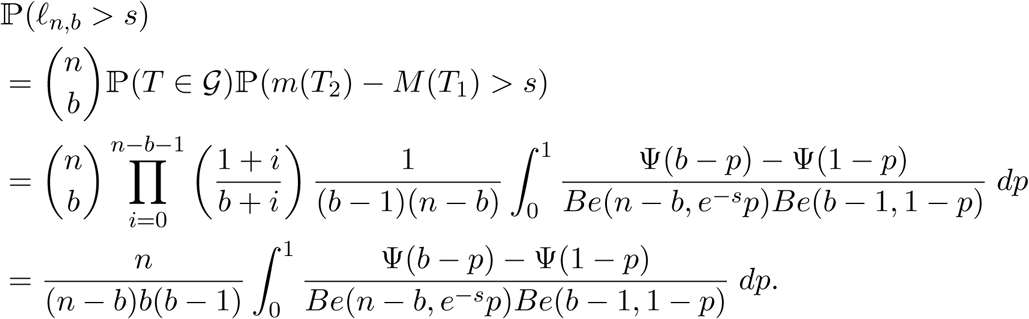

□

### Proof of Corollary 4.2.

Observe that, uniformly for *p* ∈ (0, 1), we have

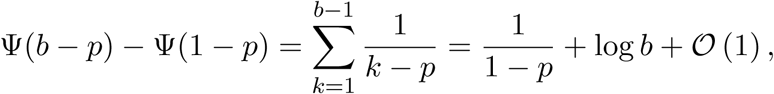

thus, substituting in (24) and also using Stirling’s approximation and Euler’s reflection formula, we obtain

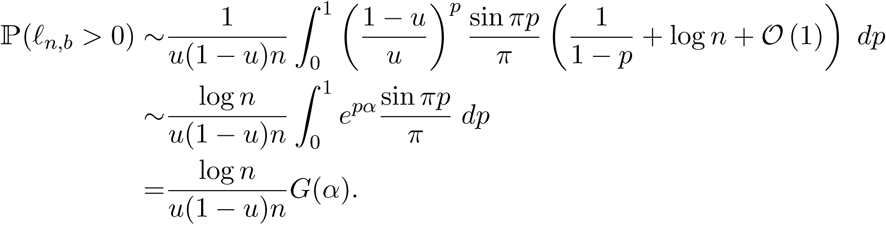

On the other hand, for any *s* > 0 we have

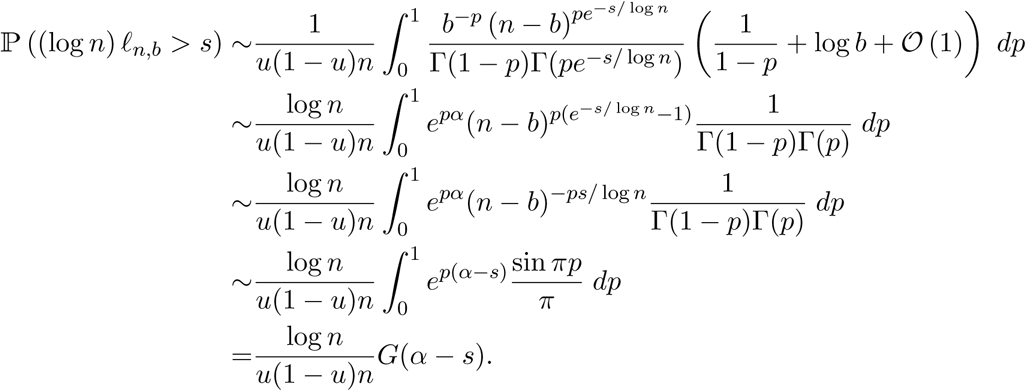

□

### Proof of Theorem 4.3.

Letting 𝓁_*π*_ := ℒ (*t* : *π* ∈ Π^*n*^(*t*)) for any subset *π* ⊂ [*n*], by exchangeability of Π^*n*^(*t*) we have

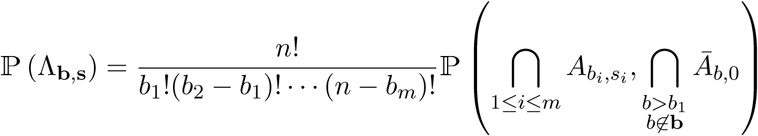

where

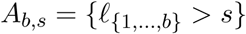

and

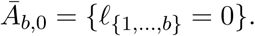

Recall that 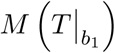 is defined as the maximum of the exponential edges associated to the root of 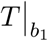. Letting *b*_*m*+1_ := *n*, and also letting *E*_*b*_, 1 ≤ *b* ≤ *n*, be the exponential variable associated to *b*, we have

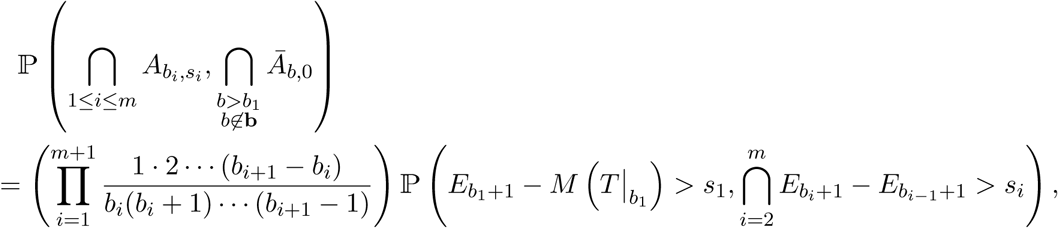

where the product above is the probability that *T* is structured in such a way that {*b*_1_ + 1} attaches to {1} and is the root of a subtree formed with {*b*_1_ + 1, …, *b*_2_}, that {*b*_2_ + 1} attaches to {1} and is the root of a subtree formed with {*b*_2_ + 1, …, *b*_3_}, and so forth. Using the independence of the exponential variables we obtain

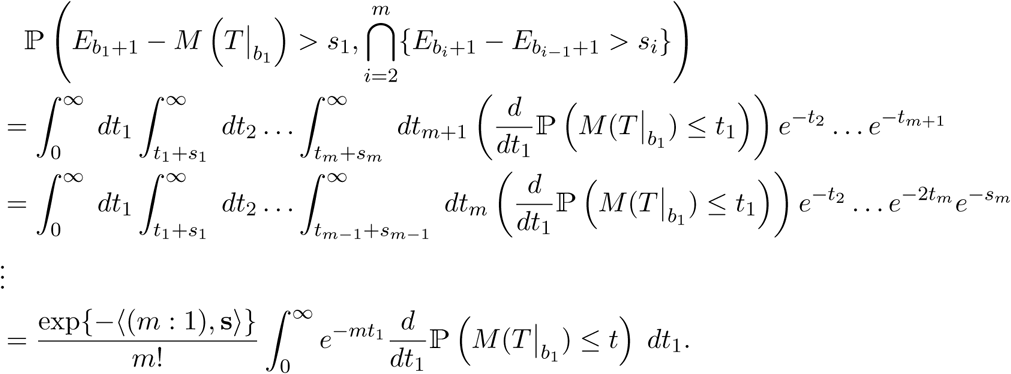

From (10) and making *p* = *e*^−*t*^ in the above integral, and putting all together we obtain (25). Finally (26) follows from

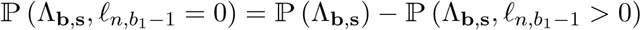

and, recursively,

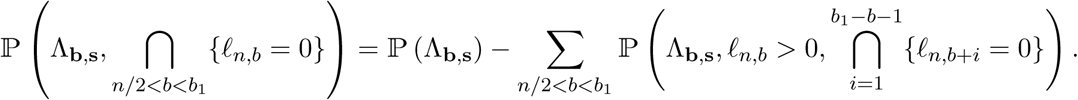

Substituting (25) in the above expression, we obtain (26). □

## Aknowledgements

Alejandro and Arno would like to thank Geronimo Uribe Bravo for his fruitful suggestions on the second moment method.

